# Selective brain cooling monitored by CT perfusion as adjuvant therapy in a porcine model of severe ischemic stroke

**DOI:** 10.1101/2022.11.11.516055

**Authors:** Olivia L.H. Tong, Kevin J. Chung, Jennifer Hadway, Laura Morrison, Lise Desjardins, Susan Tyler, Marcus Flamminio, Lynn Keenliside, Ting-Yim Lee

## Abstract

Despite the advances in ischemic stroke treatment, not all patients are eligible for or fully recovered after recanalization therapies. Therapeutic hypothermia could be adjuvant therapy that optimizes the beneficial effect of reperfusion. While conventional whole-body cooling has severe adverse effects, selective brain cooling has emerged as an attractive alternative. However, clinical application is limited by the lack of optimal delivery methods and unknown treatment parameters. Optimal parameters may depend on injury levels and monitoring cerebral perfusion may provide valuable information. Here, we show that selective brain cooling via our in-house developed Vortex tube IntraNasal Cooling Instrument (VINCI), even with a clinically relevant delay in treatment, can attenuate subacute injuries in animals with severe ischemic stroke. The treatment responses of selective brain cooling were characterized by CT Perfusion (CTP). The predicted lesion volume by CTP matched the true infarct volume by histology when the brain temperature was decreased by 5°C from normothermia. More importantly, we found that global hyperemia (high cerebral blood flow) before rewarming could be an early manifestation of poor treatment outcomes. Altogether, our study shows that VINCI-enabled brain cooling could be guided by CTP imaging as adjuvant therapy for severe ischemic stroke. This work lays the groundwork toward individualized selective brain cooling.

**Significance Statement:** Not all patients suffering from ischemic stroke are eligible or fully recovered after recanalization therapies. Therapeutic hypothermia could be an adjuvant therapy, but the clinical application is hindered by the delivery methods. The optimum treatment depth and duration are also unknown, and they may depend on the injury level. We developed a non-invasive selective brain cooling device, Vortex tube IntraNasal Cooling Instrument (VINCI). The treatment responses were characterized by CT Perfusion (CTP). Global hyperemia (high cerebral blood flow) was identified and could be an early manifestation of poor treatment outcomes. Our work shows that VINCI-enabled brain cooling could be guided by CTP imaging as adjuvant therapy for ischemic stroke. This work also lays the groundwork toward individualized selective brain cooling.

## Introduction

Despite the advances in ischemic stroke treatment, an unmet need is that not all patients are eligible for or fully recovered after recanalization therapies (1). There is a need for adjuvant therapy to extend eligibility and optimize the beneficial effect of reperfusion. Preclinical studies have shown that therapeutic hypothermia (TH) could expand treatment windows for recanalization. TH reduces tissue energy requirements to prevent infarct growth and preserve ischemic penumbra (i.e., brain tissue at risk of infarction) (2–7). However, clinical trials of whole-body cooling for ischemic stroke have been terminated early because of the serious adverse effects (8, 9). Hence, selective brain cooling has emerged as an attractive alternative. Essentially, selective brain cooling cools down the brain to a target temperature to achieve cerebroprotection, while the body is maintained at normothermia. In the past decade, different selective brain cooling methods have been investigated with variable results. Clinical applications of these methods are limited by their technical challenges. The challenges include the lack of temperature control for cold fluid infusion (10), the invasiveness of catheters for intravascular cooling (11), the requirement of extensive resources for extracorporeal blood cooling (12), and the expensive coolant of transnasal perfluorocarbon cooling (13–15). To further complicate the situation, the optimum depth and duration of hypothermia may vary depending on the type and level of injury. Therefore, clinical translation of selective brain cooling requires: (i) a non-invasive brain cooling tool that would rapidly induce and precisely maintain the required brain-body temperature gradient, and (ii) a monitoring technique that would characterize the treatment response of selective brain cooling.

For the non-invasive brain cooling device, we developed a prototype for intranasal cooling in large animals and patients, known as Vortex tube IntraNasal Cooling Instrument (VINCI) (16, 17). VINCI generates cold air directed into the nostrils using a vortex tube under the feedback control of the brain temperature. A vortex tube can decrease the temperature of the incoming air up to 20°C. Feedback control is achieved by proportional-integral-derivative controllers to automatically adjust the parameters of the vortex tube (i.e., flow rate, input pressure of compressed air, and output temperature). Selective brain cooling is not a trivial task because heat transfer between the brain and the body is facilitated by blood flow. In the present study, we increased the cooling power and improved heat retention in the body to maximize the brain-body temperature gradient.

Cerebral blood flow (CBF) plays a major role in brain temperature regulation and tissue viability. Although vital signs (i.e., physiological variables) are used to guide the management of acute brain injuries, deviations in physiological variables are delayed responses of secondary brain injuries. These parameters could not sensitively guide the duration and depth of hypothermia. Herein, we utilized computed tomography perfusion (CTP) as a monitoring technique for cerebral perfusion. CTP is an established clinical imaging technique that measures hemodynamic parameters to inform treatment decisions in routine acute stroke care (18–21).

Given the sensitivity of cerebral hemodynamics to brain temperature and physiology, we hypothesized that CTP imaging may track stroke lesion volume and characterize treatment response of selective brain cooling. We monitored the treatment response of VINCI-enabled brain cooling by CTP in a porcine model of severe transient ischemic stroke. The utility of CTP in brain cooling was evaluated by comparing the CTP results to histology. Our data show that (i) VINCI-enabled brain cooling could be adjuvant therapy of stroke, and (ii) global hyperemia (high CBF) before rewarming could be an early manifestation of poor treatment outcome.

## Results

### Characteristics of severe ischemic stroke without cooling (control)

To determine the effects of selective brain cooling, we first established a CT-guided transient severe ischemic model by intraparenchymal infusion of endothelin-1 (ET-1) via a small burr hole in the skull (*SI Appendix,* Fig. S1*A*). Our model requires minimal surgeries compared to the extensive periorbital surgery required to occlude a middle cerebral artery (22). The transient nature of ET-1 makes it ideal for evaluating reperfusion and associated hemodynamic response. More details are provided in the *Appendix, Extended Methods.*

Throughout the study, the vitals were recorded at minute intervals and scans were performed at regular intervals. Intensive care was provided for the animals for 28 ± 2 hours or until the brain was not viable. We defined the ‘brain dead’ condition when the animal was euthanized, as (i) there was global severe ischemia (CBF < 12 mL/min/100g) in the brain and (ii) the intracranial pressure was within 10 mmHg of the mean arterial blood pressure.

As a severe ischemic model, all animals without cooling (control) were brain dead 11.5 ± 2.0 h post first ET-1 injection (Fig. 1 *A and B*). At 2h, the ipsilateral CBF measured by CTP was < 75% relative to the contralateral hemisphere (Fig. 1 *F*). When blood flow was restored, there was an overshoot of blood flow (hyperemia; CBF >165%) relative to the baseline for 4 hours (Fig. 1*F*). The hyperemia was global and happened in both hemispheres (Fig. 1 *B* and *F*). This increase in blood flow relative to baseline suggested that the vasoconstricting effect of ET-1 dissipated within an hour after the last injection (Fig. 1*F*). Beginning at 7h, CBF progressively declined until the end of the study (Fig. 1*F*). Cerebral blood volume (CBV) measured by CTP had a similar response to CBF, it decreased up to 2h, then increased for 4 hours before a continuous decline until end of the study (Fig. 1*B* and *SI Appendix,* Fig. S2*A*). The steady decline of CBF and CBV mirrored the declines of blood pressure (BP) and cerebral perfusion pressure (CPP), suggesting impairment of cerebral autoregulation (Fig. 1 *C* and *F*). The ICP was greater than 20 mmHg at 4h and reached 40.9 ± 14.7 mmHg by 10 h (Fig. 1*C*). The progressive increase in ICP is consistent with malignant edema resulting from subacute injuries. These data suggested that prolonged increase in cerebral perfusion after reperfusion was associated with secondary brain injury.

**Fig. 1.**
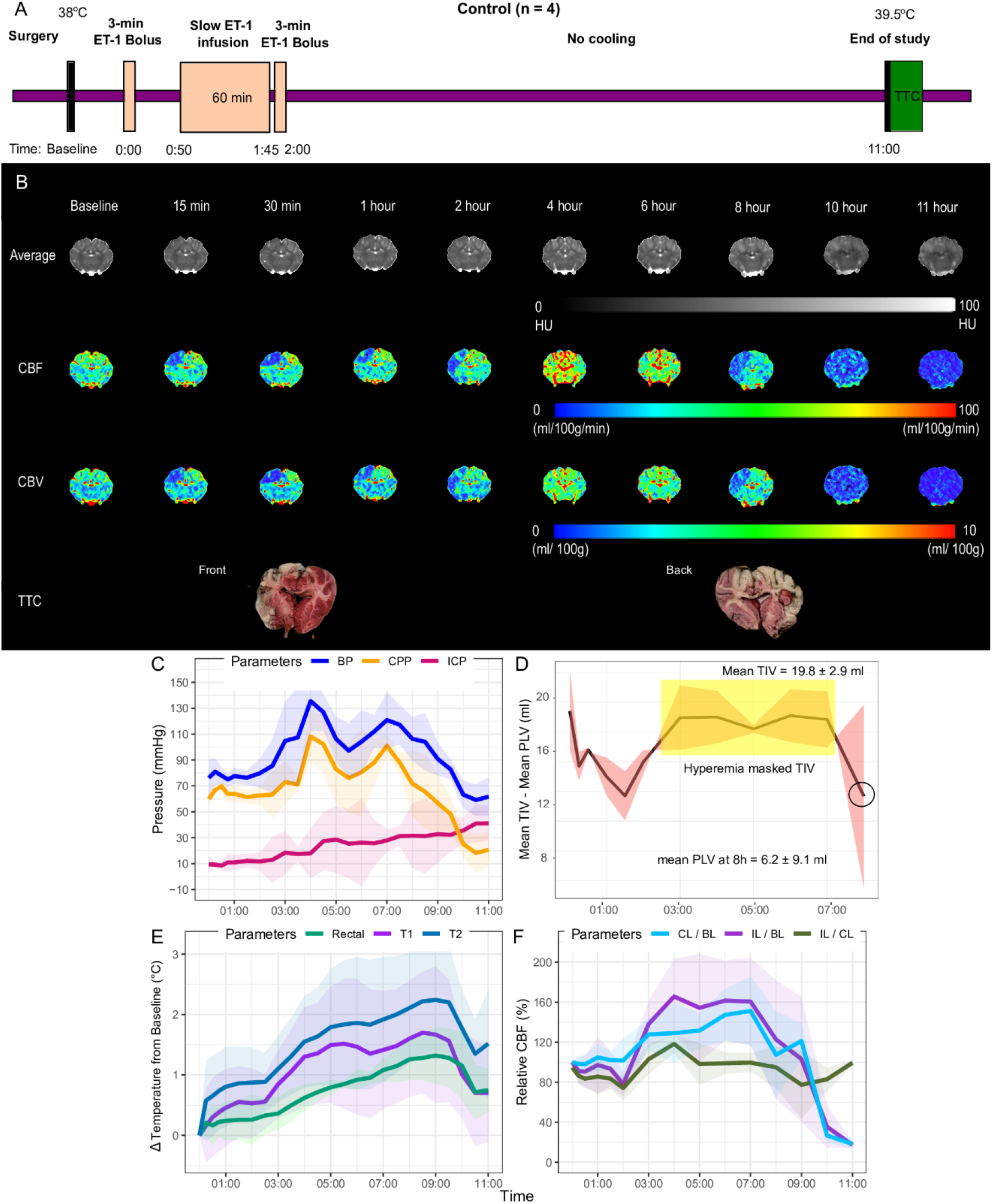
Brain hemodynamic and physiological responses in swine subject to severe ischemic stroke without cooling (i.e., control). (*A*) Scheme showing protocol of severe ischemic stroke pigs without cooling. CT-guided surgeries (burr holes) were performed on adult, female Duroc Cross pigs (n = 4). The vasoconstrictor, endothelin (ET-1; 33 μg), was dissolved in sterile water (200 μl). A catheter for infusion of ET-1 was inserted in the burr hole above the somatosensory cortex to induce ischemia temporarily. The ET-1 injection schedules were as follows: a bolus, a 50 min slow infusion after the first dose, and a bolus immediately after the completion of the second dose. Throughout the study, the vitals and CT perfusion (CTP) were recorded at regular intervals. In this group, experiments were terminated when the brain was not viable as determined by CTP (blood flow < 12 ml/min/100g in the entire brain). After the animals were euthanized, the brains were excised, and brain slices were stained in 2% Tetrazolium chloride stain (TTC). (*B*) Representative average, CBF, and CBV maps at different time points for a control subject ; as well as the corresponding TTC-stained brain slice. (*C*) Mean physiological responses, (*E*) mean temperatures, and (*F*) relative cerebral blood throughout the study (11h) in control pigs (n = 4), are reported as mean ± SD. (*D*) Difference between mean true infarct volume (TIV) and CTP predicted lesion volume (PLV) at different time points, is depicted as mean ± SD. The yellow box represents the period in which hyperemia masked TIV, and the circle indicates the mean PLV at 8h which estimated TIV more closely than PLV at other time points. T1, anterior brain temperature. T2, posterior brain temperature. IL, ipsilateral hemisphere. CL, contralateral hemisphere. BL, baseline.

The vital signs (BP, CPP, ICP, and heart rate) all exhibited a biomodal response (Fig. 1*C* and *SI Appendix* Fig. S2*D*). Brain temperatures progressively increased with peak differences of 1.7 ± 1.0°C (anterior) and 2.2 ± O.9°C (posterior) from the baseline at 8.5h (Fig. 1*E*). Similarly, the body (rectal) temperature showed an increasing trend that declined after 9h (Fig. 1*E*), indicating the development of fever after ischemic stroke.

To evaluate the utility of CTP for selective brain cooling, the mean predicted lesion volume (PLV) defined with absolute CBF-CBV threshold from CTP scans at different study time points was compared to the mean true infarct volume (TIV) determined by histology (2,3,5-Triphenyltetrazolium chloride; TTC) (Fig. 1*D*). The PLV by the final CTP scan with global hypoperfusion (CBF < 12 ml/min/100g in the entire brain) overestimated TIV (Table 1), while that from CTP at global hyperemia underestimated TIV (Fig 1 *B* and *D*). On the other hand, PLV from CTP scans at earlier time points (2h and 8h) estimated TIV more closely (Table 1). Given that the final ET-1 injection was completed 5 min before the 2h scan, these data showed that CTP tracked the initial stroke lesion volume but the infarct growth due to secondary injury was not immediately reflected. Altogether, our data characterized the response of this transient severe ischemic pig model.

**Table 1.** Infarct volumes defined by histology (TIVs) and CTP predicted lesion volumes (PLVs) at significant phases of the experiment.

| Condition | TIV by histology (mean ± SD; ml) | Phase | CTP scan | PLV by CTP (mean ± SD; ml) |
| --- | --- | --- | --- | --- |
| NC (N = 4) | 19.8 ± 2.9 | Last ET-1 injection | 2 h | 6.1 ± 4.6 |
| | | Hyperemia | 4 h | 0.9 ± 0.7 |
| | | Before global hypoperfusion | 8 h | 6.2 ± 9.1 |
| | | Final scan – global hypoperfusion | 10h | 54.0 ± 6.7 |
| TH34 (N = 4) | 21.5 ± 6.0 | Last ET-1 injection/ Induction | 2 h | 5.6 ± 3.1 |
| | | Maintenance at 34°C | 3 h | 7.8 ± 7.7 |
| | | Hyperemia | 10h | 1.8 ± 2.6 |
| | | Before global hypoperfusion | 14h | 6.2 ± 4.8 |
| | | Final – global hypoperfusion | 15h | 40.2 ± 12.7 |
| TH33 (N = 4) | 5.3 ± 2.3* | Last ET-1 injection/ Induction | 2 h | 2.5 ± 2.3 |
| | | Maintenance at 33°C | 3 h | 5.1 ± 3.6 |
| | | Rewarm | 20h | 0.6 ± 0.2 |
| | | Final scan before end of study | 28h | 0.7 ± 0.7 |

\*p < 0.05, non-parametric Kruskal Wallis tests with Bonferroni correction – TIV of TH33 was significantly different from the TIV of TH34 and NC

### VINCI-enabled selective brain cooling

An effective and reliable intranasal cooling device is desired for use in hospitals. To maximize the brain-body temperature gradient, a compact Peltier pre-cooler and an intravenous fluid warmer were added to the system. The Peltier pre-cooler increased cooling power by reducing the hospital air temperature to 10°C, and the vortex tube further cools the air to −10°C; the induction time decreased from 5 hours to < 75 minutes. The intravenous fluid warmer provided warmed fluids to improve heat retention in the body, and the body temperature was maintained within 2.1 ± 0.4°C from baseline (normothermia) throughout the study. The optimized VINCI is shown in *SI Appendix* Fig. S5 *A-C*.

Clinical studies for acute ischemic stroke reduced body temperature by 3 to 4°C (i.e., the target temperatures between 33°C and 34°C) (23, 24), so we examined the effect of a 4°C and 5°C decrease in brain temperature (TH34 and TH33 respectively) by VINCI in the severe ischemic stroke model. Cooling started at 2h after stroke onset, then the brain temperature was maintained at the target temperature for 17h, followed by rewarming at a rate of 0.5°C/ h (Fig. 2*A* and 3*A*/ The anterior brain temperature was maintained at a 33.1 ± 0.3°C and 34.3 ± 0.6°C throughout the maintenance period in TH33 and TH34 respectively, indicating VINCI maintained temperature precisely (Fig. 2*E* and 3*E*). The brain-body temperature gradient was the largest (around 4.5°C) during induction, then the average brain-body temperature gradient was maintained at 2.8 ± 0.4°C during maintenance (*SI Appendix* Fig. S4 *A-C*), suggesting selective brain cooling. Lastly, except for the initial difference during induction, the temperature difference between the anterior and posterior brain was around 0.5°C with and without cooling (*SI Appendix* Fig. S4 *A-C*).

**Fig. 2.**
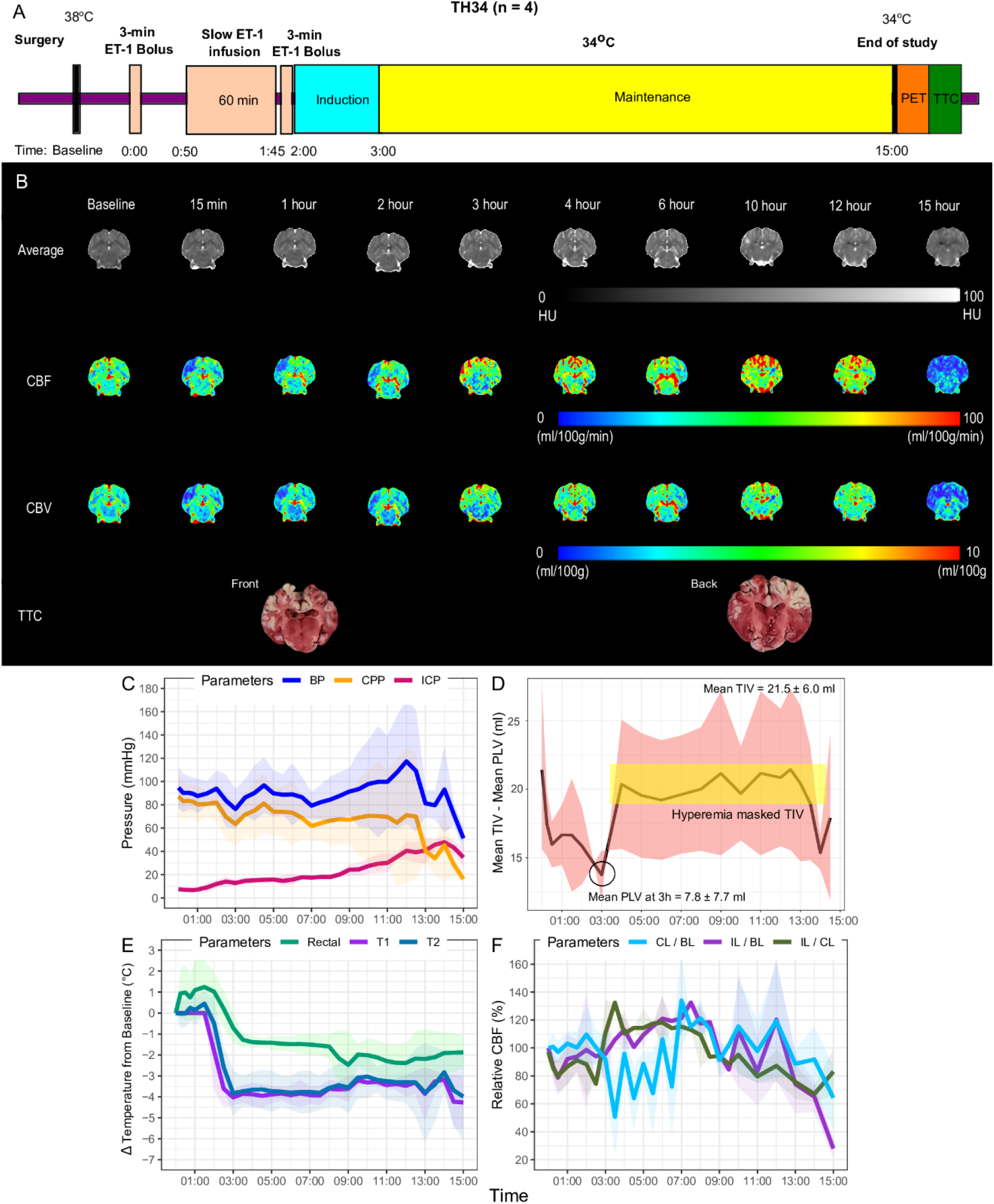
Brain hemodynamic and physiological responses in TH34 for swine subject to severe ischemic stroke. (*A*) Scheme showing protocol of severe ischemic stroke pigs in TH34. CT-guided surgeries (burr holes) were performed on adult, female Duroc Cross pigs (n = 4). Similar to the control, the ET-1 injection (33 μg of ET-1 in 200 μl of water) schedules were as follows: a bolus, a slow infusion 50 min after the first dose, and a bolus immediately after the completion of the second dose. Cooling was initiated at 2h after the first ET-1 injection, and the brain temperature reached 34°C in an hour. The temperature was maintained at 34°C until the end of the study. Throughout the study, the vitals and CT perfusion (CTP) were recorded at regular intervals. In this group, experiments were terminated when the maximum allowed anesthetic doses could not keep the animal anesthetized, or when the brain was not viable as determined by CTP (blood flow < 12 ml/min/100g in the entire brain). PET imaging was performed on one TH34 animal before sacrifice. After the animals were euthanized, the brains were excised, and brain slices were stained in 2% Tetrazolium chloride stain (TTC). (*B*) Representative average, CBF, and CBV maps at different time points for a subject in TH34; as well as the corresponding TTC-stained brain slice. (*C*) Mean physiological responses, (*E*) mean temperatures, and (*F*) relative cerebral blood throughout the study in TH34 pigs (n = 4), are reported as mean ± SD. (*D*) The difference between mean true infarct volume (TIV) and predicted lesion volume (PLV) at different time points, is depicted as mean ± SD. The yellow box represents the period in which hyperemia masked TIV, and the circle indicates the mean PLV at 3h which estimated TIV more closely than PLVs at other time points. T1, anterior brain temperature. T2, posterior brain temperature. IL, ipsilateral hemisphere. CL, contralateral hemisphere. BL, baseline.

**Fig. 3.**
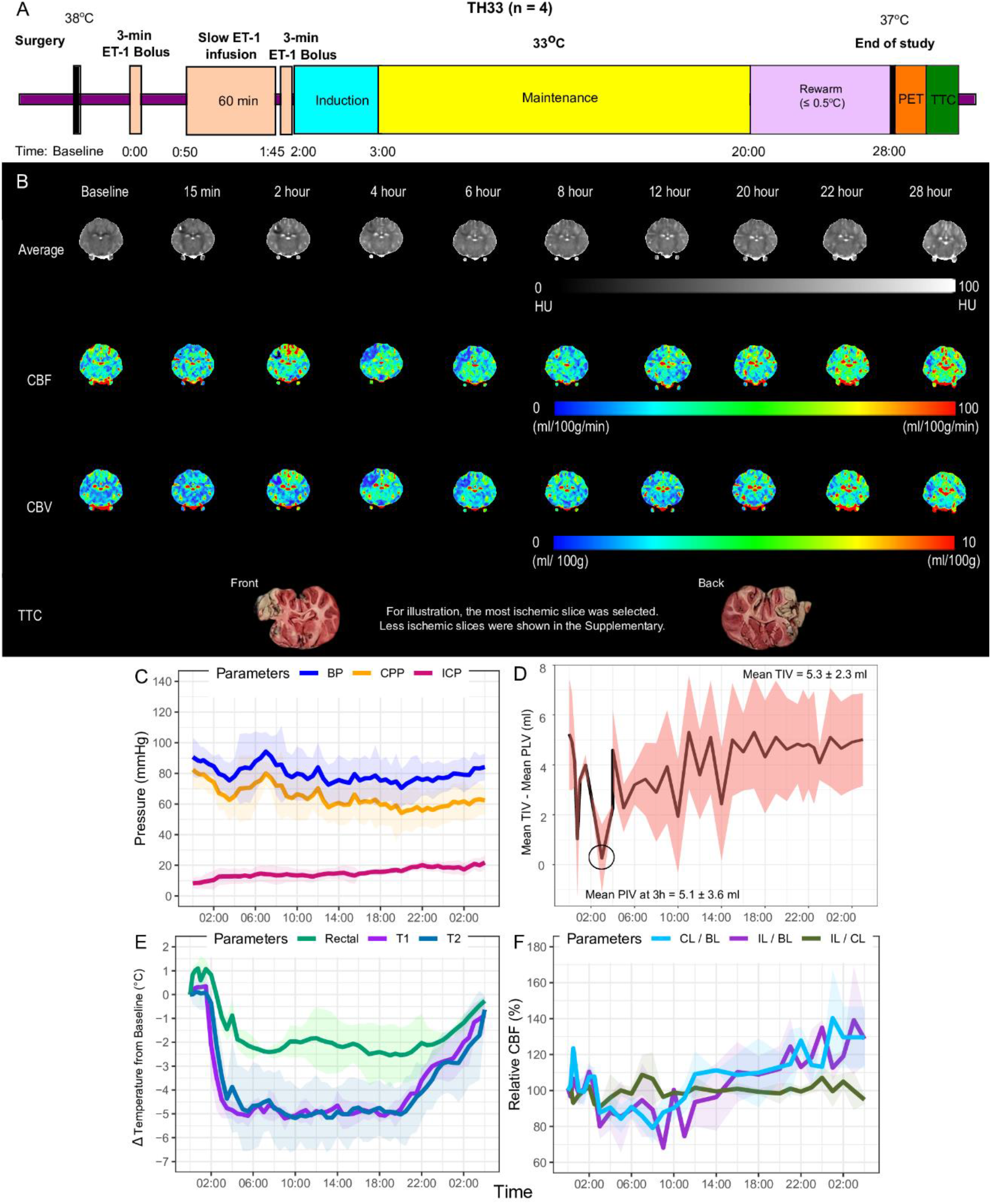
Brain hemodynamic and physiological responses in TH33 for swine subject to severe ischemic stroke. (*A*) Scheme showing protocol of severe ischemic stroke pigs in TH33. CT-guided surgeries (burr holes) were performed on adult, female Duroc Cross pigs (n = 4). Similar to the control, the ET-1 injection (33 μg of ET-1 in 200 μl of water) schedules were as follows: a bolus, a slow infusion 50 min after the first dose, and a bolus immediately after the completion of the second dose. Cooling was initiated at 2h after the first ET-1 injection, and the brain temperature reached 33°C in an hour. The temperature was maintained at 33°C for 17 hours (i.e., 20h post) before rewarming at a rate of 0.5°C /hour. Throughout the study, the vitals and CT perfusion (CTP) were recorded at regular intervals. In this group, experiments were terminated at 28 ± 2 hours. PET imaging was performed on three TH33 animals before sacrifice. After the animals were euthanized, the brains were excised, and brain slices were stained in 2% Tetrazolium chloride stain (TTC). (*B*) Representative average, CBF, and CBV maps at different time points for a subject in control; as well as the corresponding TTC-stained brain slice. (*C*) Mean physiological responses, (*E*) mean temperatures, and (*F*) relative cerebral blood throughout the study in TH33 pigs (n = 4), are reported as mean ± SD. (*D*) The difference between mean true infarct volume (TIV) and predicted lesion volume (PLV) at different time points, is depicted as mean ± SD. The yellow box represents the period in which hyperemia masked TIV, and the circle indicates the mean PLV at 3h which estimated TIV more closely than PLVs at other time points. T1, anterior brain temperature. T2, posterior brain temperature. IL, ipsilateral hemisphere. CL, contralateral hemisphere. BL, baseline.

The heart rate of TH33 during maintenance (65.4 ± 4.7 bpm) was lower than the normal range of pigs in this age group, and it was significantly lower than no cooling (*P* = 0.04) and TH34 (*P* = 0.001; *SI Appendix* Fig. S2*D*). Although we did not use an electrocardiogram to monitor the heart condition, we did not observe any irregular heartbeat on the vital signs monitor. Other complications, such as coagulopathies or acute infections, were also not observed.

### Characteristics of partial cerebroprotection by VINCI-enabled brain cooling (TH34)

Animals in TH34 were euthanized at 15.7 ± 3.6 h post; 3 out of 4 were brain dead at 17.3 ± 2.1 h before rewarming, which was longer than without cooling. The remaining one was euthanized early because the anesthetics were unable to keep the animal anesthetized. Although less severe than the control (no cooling), hyperemia was observed during maintenance in this group (Fig. 2 *B* and *F*). From 3h, ipsilateral CBFs progressively increased and peaked at 7h, followed by a progressive decline until their collapse (Fig 2*F*). Notably, global hyperemia (relative CBFs in both hemispheres > 130%) appeared at 7h (Fig. 2 *B* and *F*). Comparatively, the ICP was 20 mmHg at 7h but reached 47.9 ± 5.6 mmHg by 14h (Fig. 2*C*). The appearance of global hyperemia preceded the increase in ICP. Since hypothermia reduces tissue energy requirements, CBF is expected to decrease during the maintenance phase of selective brain cooling. Therefore, the presence of global hyperemia during maintenance was an early manifestation of inadequate cooling, and lower temperature was required to attenuate secondary brain injury.

For the vitals, there was a gradual decrease of CPP (Fig. 2*C*) between 5h to 12h, followed by a decline in the last hour. The data indicated that cerebral autoregulation was present initially (5 to 12 h post) and became impaired in the final hours. In addition, brain temperature increased from the maintenance temperature of 34°C after 7h post (Fig. 2*E*) despite increase in cooling by VINCI. This finding suggested that the inflammatory cascade was activated and supported the hypothesis that lower temperature was required to prevent subacute injuries.

For evaluation of CTP utility, there was no significant difference between the mean TIV of the control and TH34 groups (*p > 0.05*), and the PLV of TH34 at 3h provided the closest estimate of the TIV (Table 1 and Fig. 2*D*). Consistent with the findings in the control, the data in TH34 showed that CTP tracked the initial stroke lesion volume but not the infarct growth. A decrease in ^18^F-FFMZ uptake was also observed in the anterior brain, suggesting irreversible tissue damage (*SI Appendix* Fig. S6*A*). Together, our data characterized the treatment response of a partial benefit of cerebroprotection in severe ischemia by VINCI-enabled brain cooling to 34°C.

### Characteristics of attenuated subacute injuries by VINCI-enabled brain cooling (TH33)

Whereas 3 out of 4 TH34 animals were brain dead by 17.3 ± 2.1 h post, all in TH33 survived. The brain temperature was well maintained at target temperature ± 0.5°C and the body temperature also remained relatively stable during the maintenance period (Fig. 3*E*). Relative to the baseline, the ipsilateral CBF of TH33 was between 95% and 110% before rewarming at 20h (Fig. 5a and 5e) as opposed to > 130% before hemodynamic collapse for the control group, suggesting that severe hyperemia during reperfusion was suppressed. ICP during maintenance was 16.3 ± 0.9 mmHg and rewarming was 20.5 ± 1.0 mmHg (Fig. 3*C*). At the end of the experiment, TH33 had better ICP (*P* = 0.04) and CBF (*P* = 0.02) compared to the other two groups, indicating that the subacute injuries were attenuated. Throughout the study, the BP of TH33 (79.1 ± 4.9 mmHg; Fig. 3*C*) was also less varied than control (93.2 ± 20.9 mmHg; Fig 1*C*) and TH34 (89.2 ± 12.8 mmHg; Fig 2*C*). Consequently, impairment of cerebral autoregulation, which happened in the other two groups, was also prevented in TH33. These data suggested that the absence of global hyperemia during maintenance was associated with intact cerebral autoregulation and salvageable brain tissues were preserved.

While the BP and CBF were lower than the other two groups during maintenance, CBF was substantially higher during rewarming (Fig. 3 *B* and *F*). The increase was not correlated with worse outcome because ICP was well maintained throughout the study: 15.0 ± 3.3 mmHg. As well, the higher CBF during rewarming suggested it could be a mechanism to meet the needs of the expected increase in brain oxygen consumption and glucose metabolism increase (7). CBV mirrored the CBF pattern, and CBV gradually increased from 2.7 ± 0.2 ml to 3.2 ± 0.1 ml by the end of the study (*SI Appendix,* Fig. S2*C*). These results indicated that cerebral autoregulation was intact throughout the study.

TTC staining revealed that the mean TIV of TH33 was significantly smaller than the other two groups (*P* = 0.04; Table 1, *SI Appendix,* Fig. S8 *A-C*). Importantly, mean PLV by CTP at 3h matched the mean TIV in TH33 (Fig. 3*D*, Table 1) suggesting the cooling was able to limit the expansion of infarct from 3 h post, when brain temperature reached 33°C. As for ^18^F-FFMZ uptake, irreversible tissue damage on individual slice could not be accurately estimated because some planar infarct of TH33 were smaller (<0.5 ml) than the resolution of the PET/CT scanner (*SI Appendix* Fig. S6*B*). Altogether, these data supported the utility of CTP for VINCI-enabled brain cooling, and that global hyperemia before rewarming was an early manifestation of subacute injuries during hypothermia.

## Discussion

Despite the robust cerebroprotective effects demonstrated in preclinical studies, clinical application of TH is limited because existing delivery methods lead to adverse effects in patients. In the past decades, selective brain cooling has been proposed as a promising approach to ameliorate ischemic injuries while limiting serious complications (12, 13, 25). The present study shows that selective brain cooling via our in-house developed Vortex tube IntraNasal Cooling Instrument (VINCI) can attenuate subacute injuries in pigs subject to severe ischemic stroke with malignant edema. Importantly, CTP imaging can characterize treatment response of selective brain cooling, and that global hyperemia before rewarming is an early manifestation of subacute injuries.

Our in-house developed VINCI can non-invasively reduce brain temperature by 5°C in 1h, precisely maintain brain temperature for 17h, and rewarm the brain at an accurate rate of 0.5°C /h. We generated a severe ischemic stroke model by infusion of ET-1. Endothelin induced a diffused vasoconstriction around the infusion site which lasts for an hour (26). In this study, therapeutic cooling was initiated 2 hours after stroke onset (1^st^ infusion of ET-1) but within 10 minutes after the last (3^rd^) infusion. In other words, vessels were still occluded until around an hour after cooling began. Our study schedule, therefore, simulated a clinically relevant delay in treatment, where therapeutic cooling is only initiated on an ambulance on route to hospital and continued after reperfusion therapy. Furthermore, the current study showed that VINCI-enabled cooling can attenuate subacute injuries without severe adverse events in this transient ischemic stroke model. As such, VINCI is simple and safe to use. VINCI has adequate cooling power to selectively cool down the brain, while maintaining body temperature over a long period. By maximizing benefit and minimizing adverse effects, VINCI could be used as adjuvant therapy to make hypothermia trials for acute stroke feasible again.

This study incorporated CTP imaging to characterize treatment response. A unique finding of our work was prolonged global hyperemia was associated with secondary brain injury (Fig. 1*B* and 2*B*). Although transient reactive hyperemia during reperfusion was considered harmless, our results agree with other studies in which prolonged severe hyperemia was predictive of unsalvageable infarcted tissues even with successful recanalization (27–29). In the current study, global hyperemia was observed before deviations in physiological variables in the TH34 group, i.e., global hyperemia at 7h compared to the high blood pressure and ICP at 12 h post before collapse (Fig. 3*C* and 3*F*). CBF is expected to decrease during the maintenance phase of hypothermia, and any increase in cerebral perfusion during this period indicates insufficient cooling. As an additional test, we performed VINCI-enabled brain cooling in two animals with less severe ischemia, and severe global hyperemia was absent during the maintenance of 34°C in both surviving animals (i.e., ICP < 20 mmHg throughout the study; *SI Appendix* Fig. S7). This confirmed that the presence of global hyperemia before rewarming was an early manifestation of subacute injuries. Mechanistically, global hyperemia can be explained by arterio-venous (AV) shunts and cerebral autoregulation. In the ipsilateral hemisphere, reactive hyperemia was observed in the infarct regions in our CTP maps (Fig. 1*B*). The opening of AV shunts could be the underlying mechanism of our observation because AV shunts opened during hypoxemia (30, 31). In the contralateral hemisphere, hyperemia developed as blood flow passively followed blood pressure in both control and TH34 pigs (Fig. 1 *B-C* and 2 *B-C*). This contralateral hyperemia could be due to impairment of cerebral autoregulation (32, 33). Hyperemia occurred because blood pressure was beyond the upper limit of autoregulation, and blood vessels could not constrict to limit blood flow. By the end of the study, global hyperemia subsided, and blood flow fell below the viability threshold as ICP was >40 mmHg while CPP was <15 mmHg (Fig. 1*C* and 2*C*). In contrast, global hyperemia was absent throughout the induction and maintenance phase of TH33, and all animals survived in this group. The reactive hyperemia during rewarming was transient and well tolerated (ICP < 20 mmHg; Fig. 3*C*). Since blood flow was kept at a relatively constant level by changes in BP and CPP (Fig. 3 *C and F*), autoregulation in TH33 was intact. Overall, these findings suggest that global hyperemia during maintenance indicates insufficient cooling, and lower temperature was required.

To evaluate the utility of CTP for selective brain cooling, infarct volumes were automatically segmented from CTP maps and TTC staining of excised brain tissue (*SI Appendix* Fig. S6*C*, Table 1). One counter-intuitive finding was that the mean PLV defined by the final CTP scan with global hypoperfusion overestimated the mean TIV in the control and TH34 pigs. This discrepancy can be attributed to the timing of euthanasia. TTC stains tissues with viable mitochondria function red but does not stain ‘dead’ tissues (34–36). However, complete breakdown of mitochondria function does not happen immediately (34, 37). While complete recovery was possible after 17 min. of occlusion, a 60-min. occlusion with CBF ≤ 12 ml/100 gm/ min throughout the period still resulted in partial damages only (38, 39). In other words, irreversible injury to mitochondria occurs only after a prolonged period of ischemia (> 60 min). In this work, all animals were sacrificed immediately upon imaging of global hypoperfusion, and it is possible that these incomplete injuries were not detectable by TTC. As an additional test, we euthanized an animal 4h after we imaged global hypoperfusion, and all the brain slices were unstained with TTC (i.e., infarcted, *SI Appendix* Fig. S9). This confirmed that the discrepancy between PLV defined by hypoperfusion and TIV with TTC staining arose from not enough time was allowed for ischemia to mature into infarction.

It was hypothesized that CTP imaging may track stroke lesion volume during cooling. We found that the mean PLV by CTP consistently underestimated the mean TIV by TTC in control and TH34 (Table 1, Fig 1*D* and 2*D*). In other words, CTP tracked the initial stroke lesion volume but not the infarct growth. Since these two groups represent incomplete recovery despite successful recanalization, it was likely that the infarct continued to expand between earlier hours and euthanasia to obtain post-mortem specimens. On the other hand, the mean PLV by CTP at 3h matched the mean TIV in TH33 (Table 1, Fig 3*D*). This implied that 1) expansion of infarct was prevented once the brain temperature reached the target temperature and 2) rewarming at a rate of 0.5°C/ hour did not exacerbate the damage. Altogether, these data support the utility of CTP for selective brain cooling. Importantly, the present study raises the possibility to deliver individualized VINCI-enabled brain cooling. In the future, with the advancement of mobile CT (40) capable of CTP studies in neurointensive care units, global hyperemia as determined by CTP could be used to guide the depth and duration of hypothermia in stroke.

Based on our results, VINCI-enabled brain cooling could be guided by CTP imaging as adjuvant therapy for severe ischemic stroke. Future work would determine the CBF thresholds to guide selective brain cooling for individualized treatment. Subsequent experiments will also test the previously reported non-invasive brain temperature sensors (41) to facilitate the use of VINCI in pre-hospital and in-hospital settings. This work lays the groundwork toward individualized selective brain cooling, which will improve patient care and reduce the financial burden of health care systems.

## Methods

Additional details about the methods we used in this study are provided in SI Appendix, Methods.

### Instrumentation

The Vortex tube IntraNasal Cooling Instrument (VINCI) is an automatic system that operates under the feedback control of brain temperature. It comprises of a cooling module, a control module, and an animal interface. More details are provided in the *SI Appendix, Extended Methods.*

### Animal preparation and monitoring

All animal experiments were conducted in accordance with the guidelines of the Canadian Council on Animal Care and approved by the Animal Use Subcommittee at Western University (Protocol #2017-037 and #2021-112). More details are provided in the *SI Appendix, Extended Methods.*

### Functional imaging protocol

Brain hemodynamics were monitored by CT Perfusion (CTP). Positron emission tomography (PET) imaging was also performed on some animals at the end of hypothermia treatment to predict the final infarct. More details are provided in the *SI Appendix, Extended Methods.*

### Histology

Triphenyltetrazolium chloride (TTC) histology was performed to determine the extent of dead tissue, the true infarct volume (TIV). More details are provided in the *SI Appendix, Extended Methods.*

### Statistics

Data are depicted as mean ± SD. Statistical significance was determined using non-parametric Kruskal Wallis tests with post hoc analysis. More details are provided in the *SI Appendix, Extended Methods.*

## Acknowledgments

The authors would like to thank Lihai Yu at the cyclotron at St. Joseph’s Hospital for producing the 18^F^-FFMZ for PET imaging. This study was supported by the Heart & Stroke Foundation of Canada, the Canada Foundation for Innovation (CFI), and the Ontario Research Fund (ORF).

