## Supplementary information appendix for "Selective brain cooling monitored by CT perfusion as adjuvant therapy in a porcine model of severe ischemic stroke"

**This PDF file includes:**

Supporting text
Figures S1 to Sx
SI References

**Supporting Information Text**

### Supporting Information Text

#### Extended methods

##### Instrumentation

The Vortex tube IntraNasal Cooling Instrument (VINCI) is an automatic device that operates under the feedback control of brain temperature. It comprises of a cooling module, a control module, and an animal interface. The cooling module consists of a commercially available Peltier cooler (Thermoelectric cooler, Hozee Ltd, China) and a Ranque-Hilsch Vortex Tube (Frost Free Cold Air Gun, ITW Vortec Ltd, OH, USA). The control module consists of two Arduino-based proportional-integral-derivative controllers (PID controllers) that automatically regulate the inlet compressed air pressure and fraction ratio of cold to inlet air flow rate to control the cold air temperature and a graphical user interface (GUI) to set the cooling parameters – brain target temperature and the period during which this target temperature is maintained and the rewarm rate. The animal (pig) interface consists of two nasal cannulae to pipe cold air into the nasal cavity via the nostrils, rectal and brain temperature, and intracranial pressure probes.

In these experiments, the source of compressed air was hospital medical air. The Peltier cooler decreased the compressed air to around 10$^{o}$C before input into the vortex tube. The vortex tube separated the compressed air into hot and cold streams according to the fraction ratio set by an adjustable throttle valve. From energy conservation, the temperature differential between the cold and hot air stream was a trade-off between flow rate as a function of inlet air pressure and fraction ratio. The PID controllers regulated these variables to reach the desired cold air temperature via the throttle valve of the vortex tube and an electronic pressure regulator (Pulstronic II Series 605, Numatics Inc, Highland, MI, USA). While one PID controller was used in the induction phase to generate the coldest air for the brain to rapidly reach the target temperature, both PID controllers were used to control the air temperature during maintenance and rewarming of the brain temperature. The brain temperature as the feedback control of cold air temperature was measured by a thermocouple probe inserted into the brain. A thermistor was also placed inside one nasal cannula to measure the temperature inside the nasal cavity.

In this work, hospital medical air at a pressure of 50 PSI was the source of compressed air, and the coldest output air was -10$^{o}$C at a flow rate of 70 standard litres per minute. The nasal cannulae were inserted 8 cm into the nostrils, and the average temperature in the nostrils was 7.3 ± 3.1$^{o}$C warmer than the cold air generated by VINCI. The average cooling rate was 5.8 ± 3.6$^{o}$C/ hour.

##### Animal preparation and monitoring

All animal experiments were conducted in accordance with the guidelines of the Canadian Council on Animal Care and approved by the Animal Use Subcommittee at Western University (Protocol #2017-037 and #2021-112). Adult, female Duroc Cross pigs (n = 20, weight = 33.5 ± 9.9 kg) were used for all experiments. Anesthesia was induced by masking with 3-4% isoflurane, catheters were inserted into cephalic veins for infusion of propofol (4 mg/kg) to maintain anesthesia, followed by intubation and mechanical ventilation with an oxygen-medical (2:1) air mixture for the duration of the experiment. Both femoral arteries were catheterized to monitor vitals (SurgiVet Advisor Vital Signs Monitor, Smiths Medical, MN, USA) and collect hourly blood samples to monitor blood gases and chemistry and glucose levels. In this study, breathing rate and stroke volume were adjusted to maintain the end-tidal $CO_2$ tension ($P_eCO_2$) level at normocapnia (between 37 and 42 mmHg). 2 ml of 25% dextrose solution was administered when the blood glucose was below 4.5 mmol/L. Pigs were placed in the prone position on the bed of a clinical CT scanner (Resolution 256 slice CT scanner, GE Healthcare, WI, USA), wrapped with hot water and sheet blankets. The nasal cannulae of VINCI was coated with 2% lidocaine gel for local anesthesia and inserted into the nostrils of the pigs.

A non-contrast head CT (NCCT) coronal scan (120kV, 500 mA, 1 s/rotation, matrix size = 512 x 512, slice thickness = 1.25 mm, FOV = 22 cm) was performed to locate the positions for the implanted probes. CT scanning was used to guide the reproducible placement of intraparenchymal probes and catheter. Three 1 mm burr holes were drilled through the skull of the left hemisphere with a Dremel tool. The anterior temperature, intracranial pressure, and posterior temperature probes were inserted into the holes, which were 1.5 cm anterior, 0.1 cm lateral, and 1.5 cm posterior to the bregma, respectively (*SI Appendix* Fig. S1*A*). On the right hemisphere, a 1 mm burr hole above the somatosensory cortex (*SI Appendix* Fig. S1*A*) was drilled to insert a catheter for infusion of a vasoconstrictor, endothelin-1 (ET-1, Sigma Chemical Company, MO, USA), to induce ischemia temporarily. The hole was at the CT slice where the pituitary gland was visualized. ET-1 (33 µg) was dissolved in sterile water (200 µl) and delivered by a syringe pump (NE-100, New Era Pump Systems Inc., NY, USA). As a result, the focal ischemia was targeted at the somatosensory cortex, resembling a clinical parietal stroke with distal middle cerebral artery occlusion. Different combinations of bolus (60 µl/ min) and slow infusion (3.33 µl/ min) of ET-1 were tested to establish the severe ischemic stroke model. The selected injection schedules for the experimental groups were as follows: a bolus, a slow infusion 50 min after the first dose, and a bolus immediately after the completion of the second dose.

Throughout the study, the vitals were recorded at minute intervals. The monitored vitals were heart rate (HR), blood pressure (BP), intracranial pressure (ICP), respiratory rate (RR), arterial oxygen saturation ($S_aO_2$), rectal temperature, anterior and posterior brain temperatures. Intravenous (IV) fluid was infused at 4 ml/kg/h to replenish fluid lost. During maintenance and rewarming, warm IV fluid at 42°C was infused to warm the body. Throughout the experiment, the pigs were anesthetized with intravenous infusion of propofol (10 mg/ml at 4 – 24 mg/kg/h) and fentanyl (0.5 mg/ml at 0.3-0.5 mg/kg/h). The anesthetic infusion rate was adjusted according to the vital signs and anal tone to ensure adequate anesthesia. For all animals, the experiment continued after the last ET-1 infusion until one of the following conditions was satisfied: (1) 28 ± 2 hours, (2) when the maximum allowed anesthetic doses were unable to keep the animal anesthetized, or (3) when the brain was not viable as determined by CT perfusion (CTP; blood flow < 12 ml/min/100g in the entire brain). The animals were then euthanized by intravenous infusion of potassium chloride (1-2 ml/kg, 2 mEq/ml).

*Establishment of severe stroke model.*

Existing ischemic models involved extensive periorbital surgery to occlude a middle cerebral artery (1), but minimal surgeries to prevent complications that could confound selective brain cooling was desired. Hence, our severe stroke model needed to satisfy three criteria: 1) minimal surgeries; 2) ischemia with blood flow < 12 ml/min/100g (2) that lasted > 2h in the anterior brain; and 3) an infarct core with subacute injuries that led to brain death within 20 hours without cooling. To establish such a model, four ET-1 injection schedules were investigated in groups of two pigs per schedule: (a) schedule 1 – four boluses (60 µl/ min), (b) schedule 2 - bolus and slow infusion (3.33 µl/ min), (c) schedule 3 – bolus, slow infusion, slow infusion, and bolus, and (d) schedule 4 – bolus, slow infusion after 50 min, and bolus after 60 min. Each injection was 60 min apart for the first three schedules. Cooling was not applied during stroke model development. Intracranial pressure (ICP) was less than 20 mmHg throughout the study for the first three injection schedules. Representative hemodynamic responses and histological staining for the first three schedules are shown in *SI Appendix* Fig. S1*B*. In schedule 4, the animal succumbed to subacute injuries with ICP > 40 at 10h ± 1h (Fig. 1*C*). Hence, schedule 4 was selected for this study.

*Experimental groups.*

In this work, 8 pigs were used to establish and characterize a severe ischemic stroke model. After the model was established, 12 pigs were evenly distributed into two treatment and one control groups. The treatment groups were hypothermia at 34°C (TH34; 4°C from normothermia) and hypothermia at 33°C (TH33; 5°C from normothermia). Both groups had cooling initiated at 2h after the first ET-1 injection, then maintained at the target temperature for 17 hours before rewarming at a rate of 0.5°C /hour.

#### Functional imaging protocol

*CT Perfusion imaging.*

In addition to monitoring the physiological vitals, brain hemodynamics were monitored by CT Perfusion (CTP). The imaging parameters were as follows: 80 kV, 150 mA, 1 s/rotation, matrix size = 512 x 512, slice thickness = 2.5 mm, FOV = 22 cm. Perfusion imaging consisted of injecting a bolus (4 - 5 ml/s) of non-ionic, iodinated contrast (Isovue®-370, Bracco Diagnostic, ON, Canada; 370 mg iodine/ ml) based on the animal weight (0.9 ml/kg), followed by another bolus of 0.45 ml/kg saline at the beginning of the CTP acquisition. The CTP was acquired at 1.0 s image intervals for 45 s, and then once every 15 s for a total of 150 s. CTP scans were performed at baseline (before ET-1 injection), 15 min post-ET-1, every 30 min until 2h, every 2 h until 10 h, thereafter every 4 h, and a final scan before the end of the study at 19-20 h post. To monitor tissue perfusion, CTP maps, except the ones collected between 14-18 h post (overnight), were calculated within 10 min of acquisition.

*$^{18}$F-FFMZ positron emission tomography imaging.*

Positron emission tomography (PET) imaging was also performed on some animals at the end of hypothermia treatment using a clinical PET/CT scanner (Discovery VCT, GE Healthcare, WI, USA) to predict the final infarct. The PET dynamic imaging acquisition protocol was as follows: 10 frames of 10 s, 5 frames of 20 s, 4 frames of 40 s, 4 frames of 1 min, and 18 frames of 3 min for a total duration of 64 min. A dose of the flumazenil analogue, 2'-[$^{18}$F] fluoroflumazenil ($^{18}$[F]FFMZ; 551.6 ± 139.0 MBq), was injected via the ear vein at the same time as the start of imaging. Flumazenil (FMZ) selectively binds to the γ-aminobutyric acid type A ($GABA_A$) receptor, an inhibitory receptor located on viable neurons, therefore the lack of $^{18}$[F]FFMZ binding reflects infarction. Before the PET study, a CT scan was acquired for attenuation correction. In this work, 4 treated pigs (3 pigs in TH33, and 1 pig in TH34) were imaged.

#### Image analysis

*Registration of NCCT to pig brain atlas.*

For this purpose, an MRI-based pig brain atlas(3) was registered to the NCCT. In brief, skull-striped CT images were segmented by thresholding between -5 and 150 HU and selecting connected components corresponding to the brain. Extracranial structures eliminated by morphological erosion and dilation operations and holes filled by an iterative filling algorithm. Registration of the MR images in the brain atlas to the skull-stripped CT images was initialized by a rigid and affine transformation (4), followed by a non-rigid deformable registration (5) to align brain structures locally. The brain atlas segmentation was then registered to the CT image using the non-rigid transformation determined from the previous step before resampling to CT image resolution.

*CTP maps.*

A Johnson-Wilson-Lee model-based deconvolution algorithm(6) developed in our lab was used to process CTP studies. Arterial input function was obtained from the contralateral common carotid artery. CTP parameters include cerebral blood flow (CBF), cerebral blood volume (CBV), and permeability surface area product (PS) were used to monitor tissue perfusion as the hypothermia treatment progresses.

The digitally segmented pig brain atlas was co-registered to the average map of each CTP study as described above. Pixels with a CBF > 300 ml/min/100g, CBV > 7 ml/100g, or PS >10 ml/100g/min were excluded to remove the influence of large vessels. The brain regions were grouped into ipsilateral and contralateral hemispheres using the brain atlas to obtain the corresponding CTP values. To assess the role of perfusion imaging in selective brain cooling, absolute CTP thresholds were used to predict and segment infarct core volumes at different study time points. The absolute thresholds were CBF < 12 ml/min/100g (2) and CBV < 2 ml /100g (7, 8). The predicted lesion volumes (PLV) based on absolute thresholds were then compared with the true infarct volumes (TIV) from the corresponding histological slices.

*PET image analysis.*

The [$^{18}$F] FFMZ images were registered with the NCCT head images to visually assess its uptake in the cerebral hemispheres.

#### Histology

Triphenyltetrazolium chloride (TTC) histology was performed to determine the extent of dead tissue, the true infarct volume (TIV). After the animals were euthanized, the brain was excised and cut into coronal sections (slice thickness = 5 mm). Each slice was stained in 2% solution of TTC and fixed in 10% formalin (9). Digital images of the brain slices and ruler were photographed.

Infarct volumes were automatically segmented by an in-house Fiji-based macro (ImageJ, GPL licenses) (10). Briefly, a calibration factor was first derived from the imaged ruler. Otsu's threshold and colour thresholding with saturation between 80 to 255 were performed to segment the image. The segmented areas were then summed to give the infarct volumes.

#### Statistics

Average physiological variables, CTP parameters, predicted and true infarct volumes between the three groups were compared. Omnibus tests were performed with non-parameteric Kruskal Wallis tests. Bonferroni correction for multiple comparisons was applied to post-hoc test to detect the difference between the groups. All statistical analysis was performed on R Statistical Software (version 4.1.2; R Foundation for Statistical Computing, Vienna, Austria).

**S1**

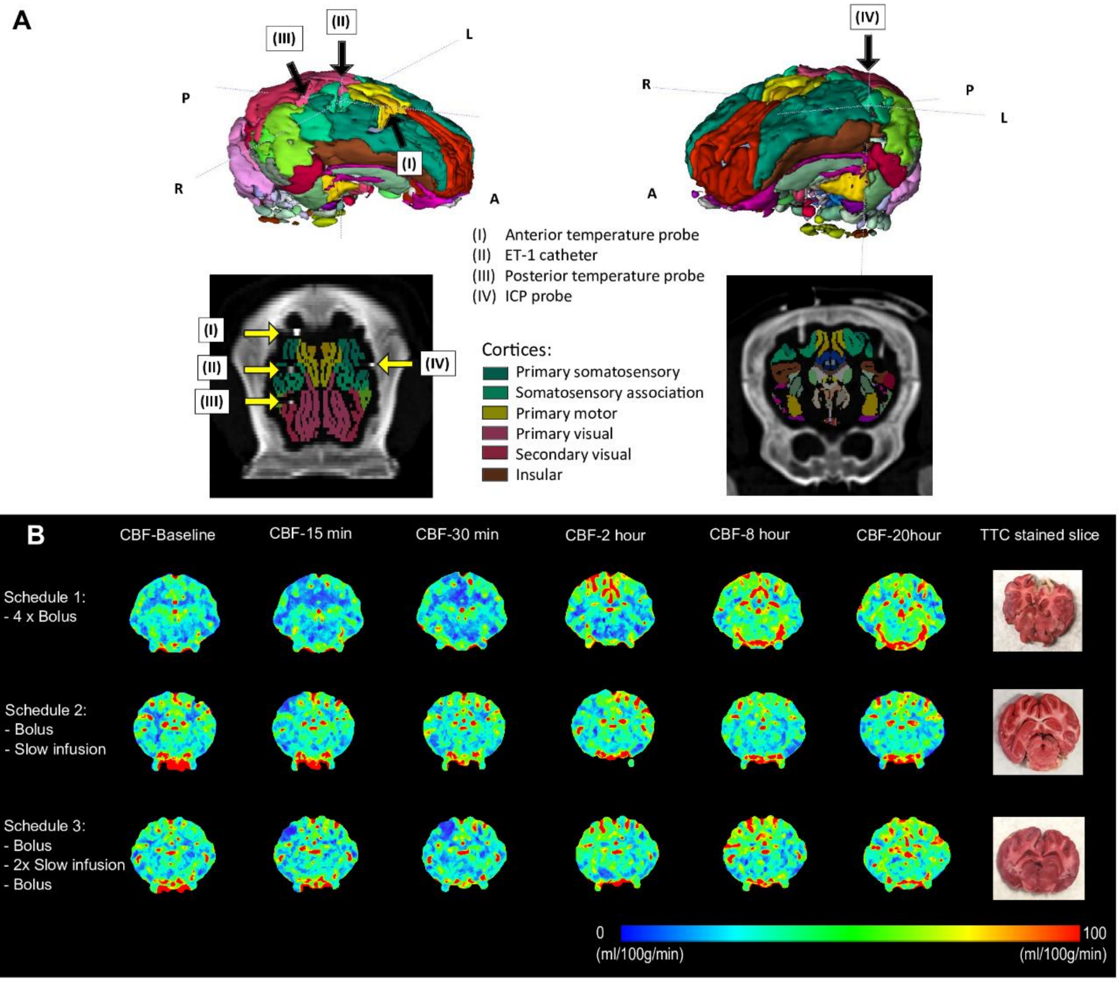


**Fig. S1.** Establishment of CT-guided porcine model of severe ischemic stroke. (*A)* Locations of implanted probes. 3D surface rendered of the pig brain (*top*) labelled with locations of implanted probes. Axial slice (*bottom left*) and coronal (*bottom right*) slice of head CT scans that showed the locations of the implanted probes and the corresponding registered brain atlas by Saikali et al.(3). (*B)* Representative cerebral blood flow maps and histological staining of different endothelin-1 injection schedules. ET-1 endothelin-1, CBF cerebral blood flow, TTC tetrazolium chloride.

**S2**

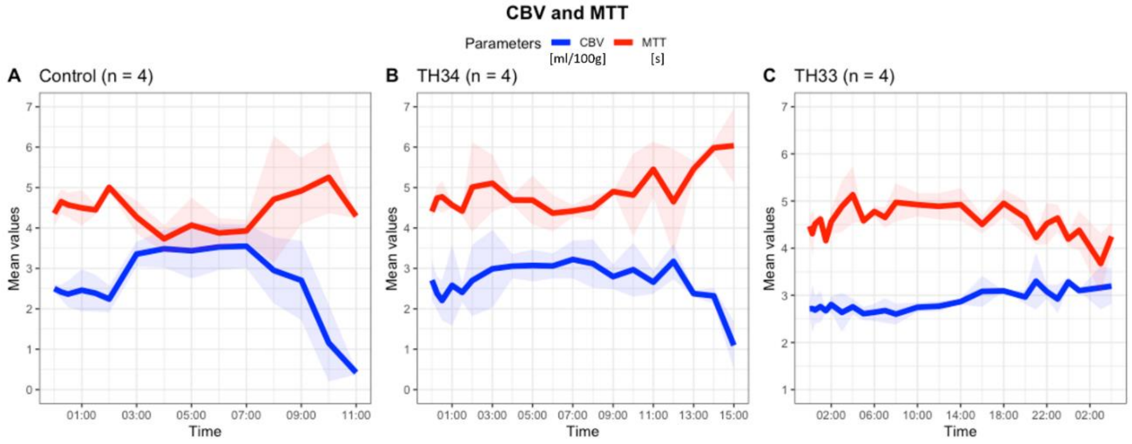


D

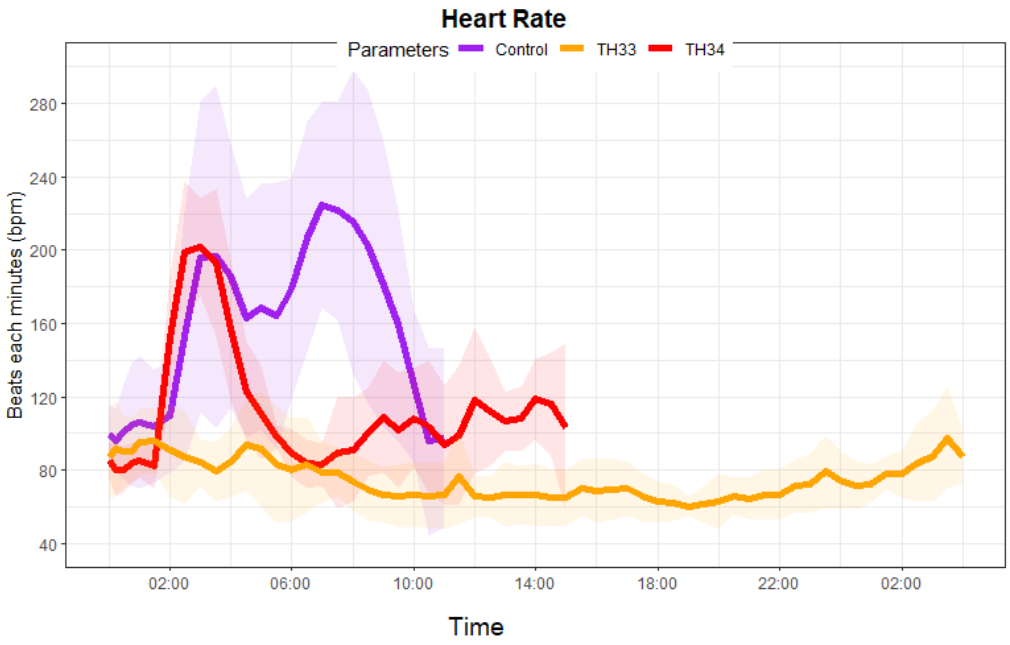


**Fig. S2**. Brain hemodynamic and physiological responses of different groups. (*A-C)* Cerebral blood volume (CBV) and mean transit time (MTT) throughout the study in control (n=4), TH34 (n=4), and TH33 (n=4) pigs, are reported as mean ± SD. (*D)* Heart rate of pigs in different groups throughout the study is depicted as mean ± SD.

**S3**

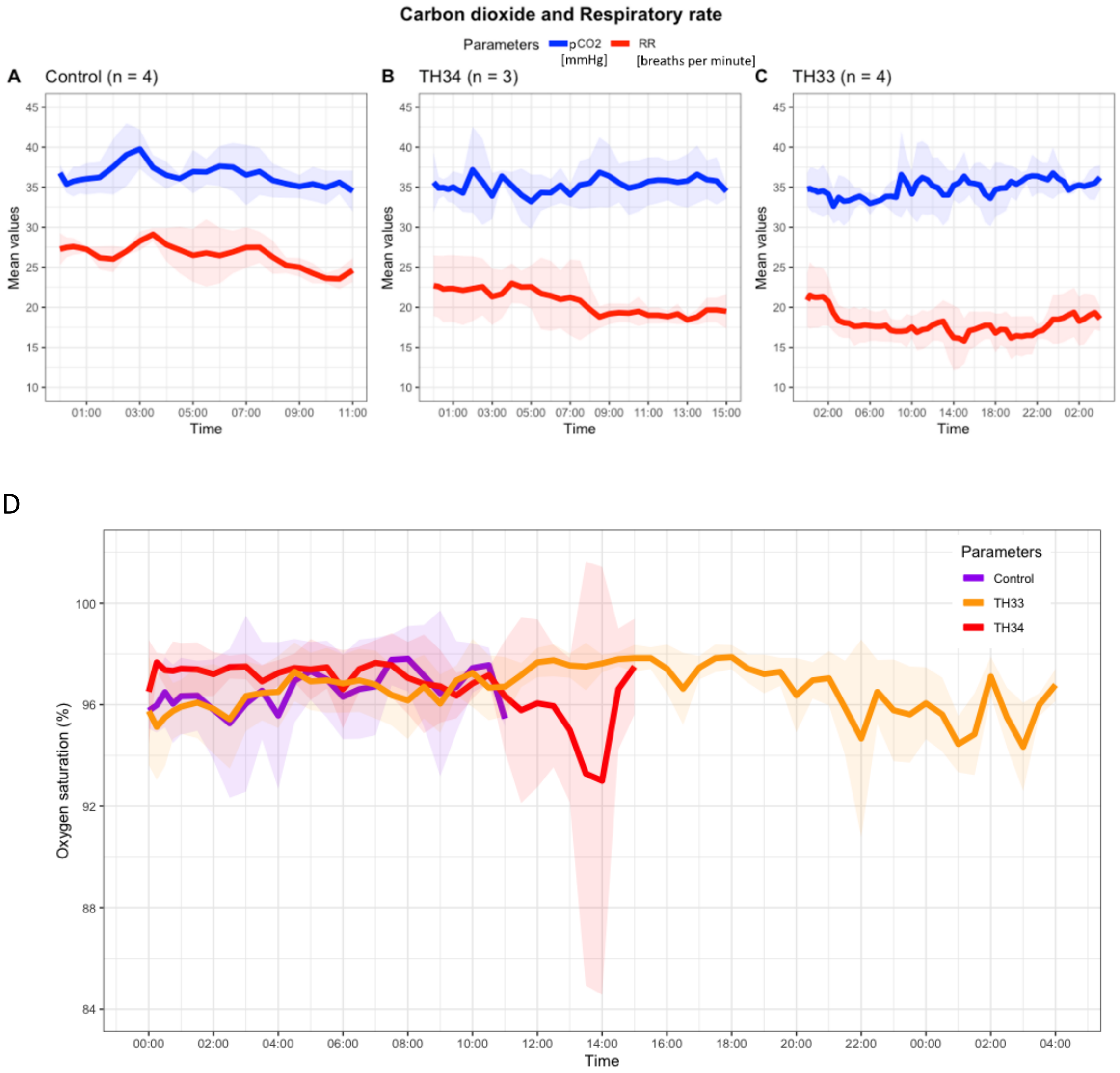


**Fig. S3**. Physiological responses of different groups. (*A-C)* Respiratory rate (RR) and partial pressure of carbon dioxide ($pCO_2$) throughout the study in control (n=4), TH34 (n=4), and TH33 (n=4) pigs, are reported as mean ± SD. (*D)* Arterial oxygen saturation ($S_aO_2$) of pigs in different groups throughout the study is depicted as mean ± SD.

## S4

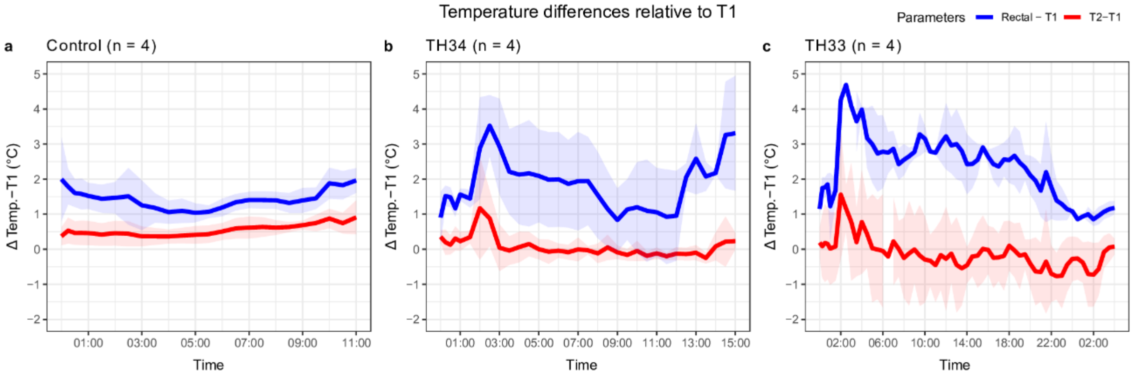


**Fig. S4.** Temperature differences relative to T1. Temperature differences in *(A)* control (n=4), *(B)* TH34 (n=4), *(C)* TH33 (n=4) pigs are reported as mean ± SD. Rectal, rectal (body) temperature. T1, anterior brain temperature. T2, posterior brain temperature. Temp., temperature.

**S5**

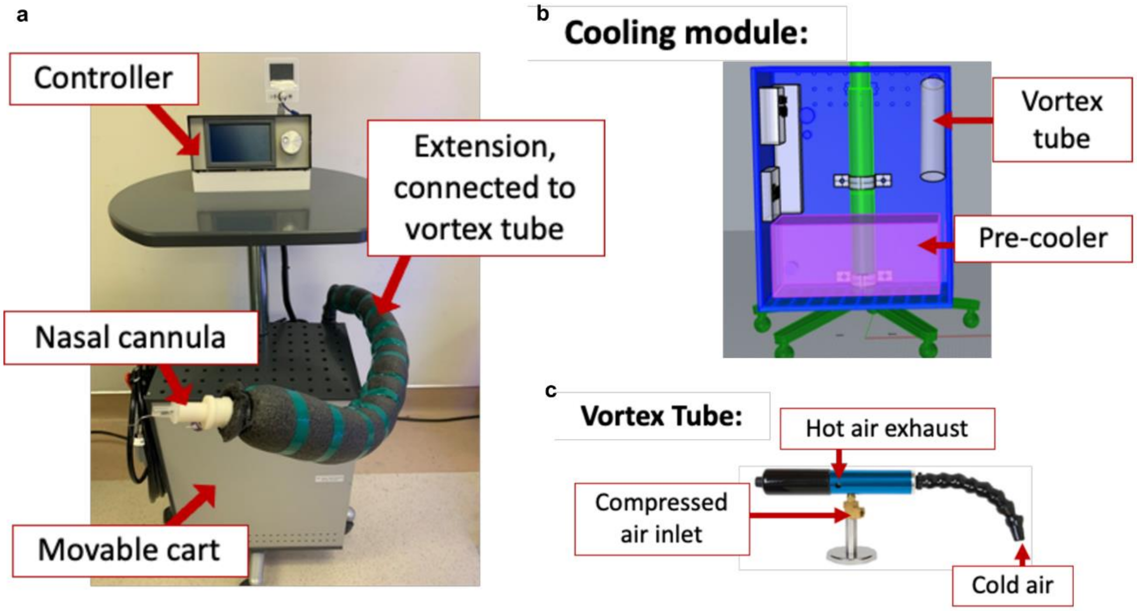


**Fig. S5.** Representations and photographs of VINCI. (*A)* Photograph of VINCI. (*B*) Cooling module of the system. (*C*) Close-up photograph of the vortex tube.

## S6

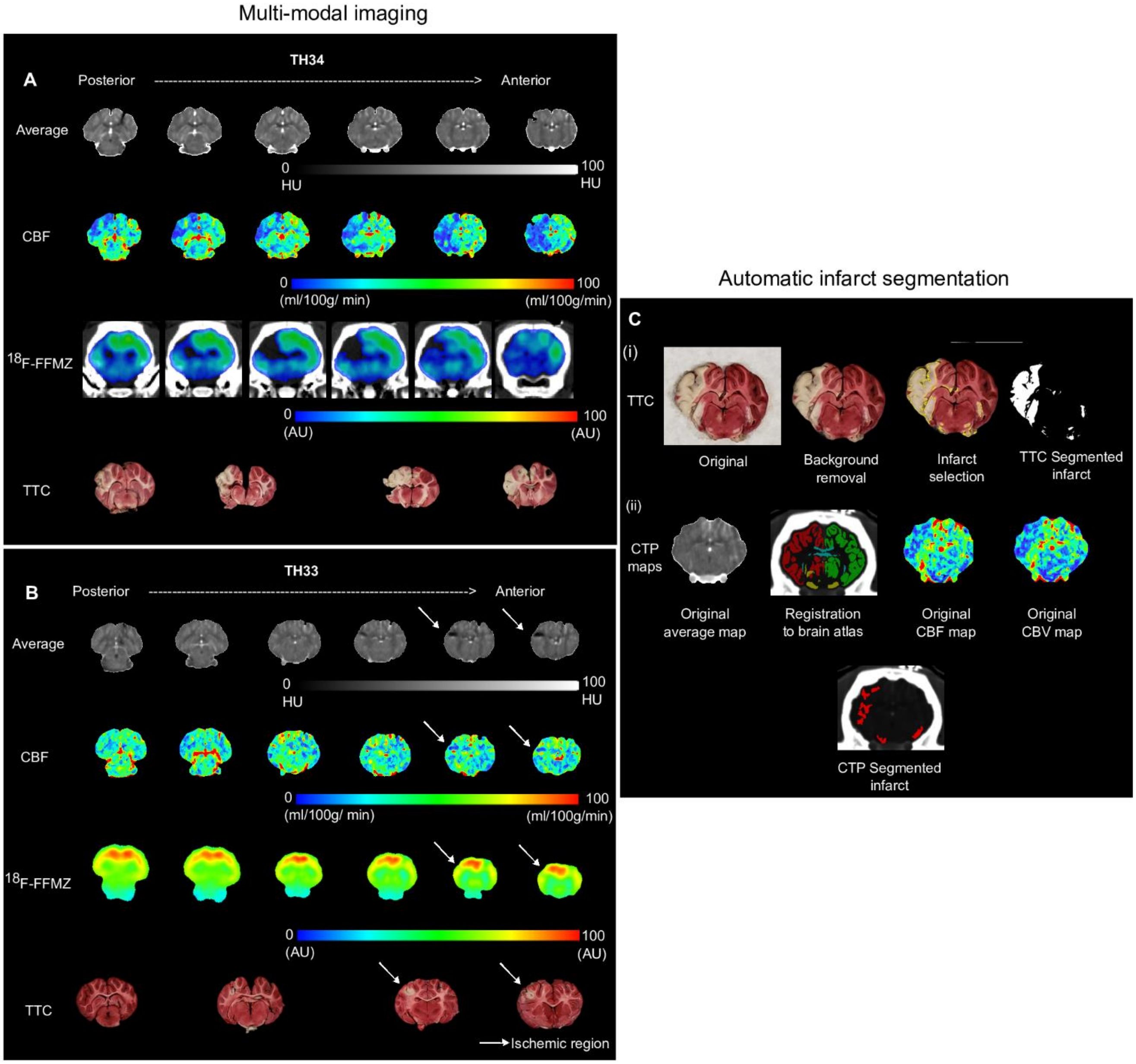


**Fig. S6.** Muti-modal imaging and infarct segmentation. Example of multi-modal imaging and TTC-stained brain in (*A*) TH34 and (*B*) TH33 pigs. (*C*) Representative automatic infarct segmentation of (*i*) TTC-stained brain slices and (*ii*) CTP maps that involved registration to brain atlas. Arrow indicates ischemic region in TTC slices and corresponding CT averge and CBF maps and PET images. $^{18}$F-FFMZ, 2'-[18F]-labelled fluoroflumazenil radiotracer for PET imaging. AU, arbitrary units.

**S7**

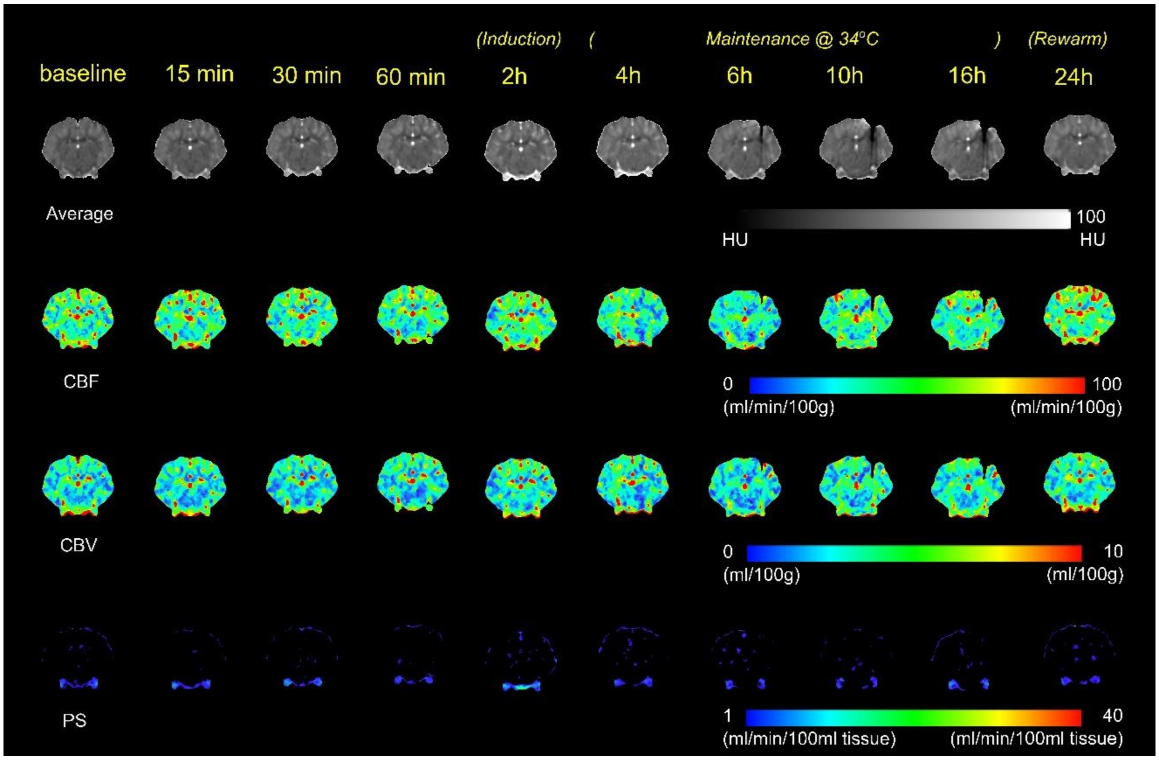


**Fig. S7.** Representative average, cerebral blood flow (CBF), cerebral blood volume (CBV), and permeability surface area product (PS) maps at different time points in TH34 swine subject to less severe ischemia. The pig survived and maintained intracranial pressure < 20 mmHg throughout the study.

**S8**

A. Control

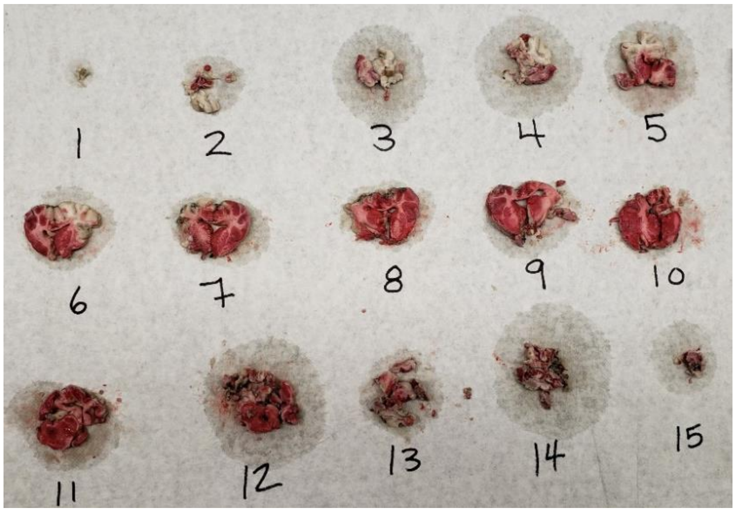


B. TH34

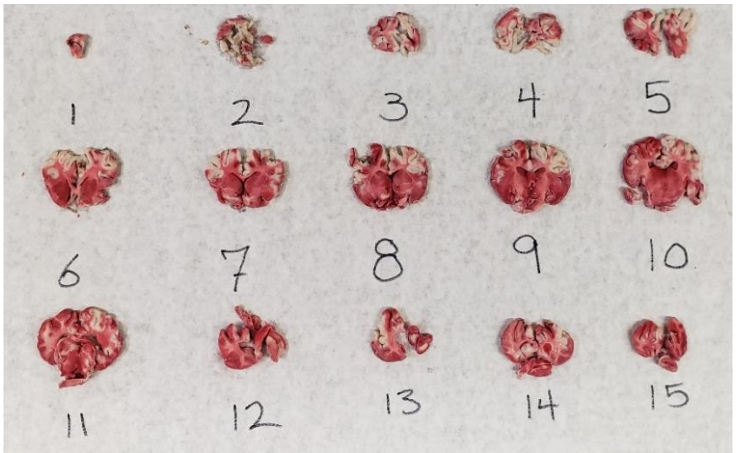


C. TH33

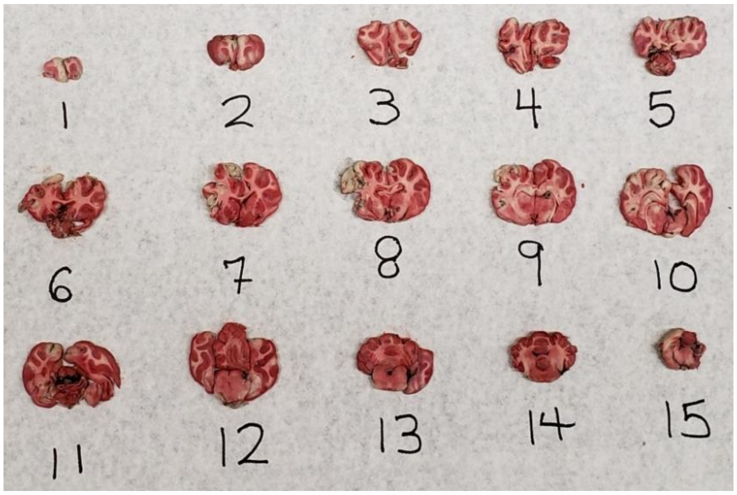


**Fig. S8.** Examples of tetrazolium chloride (TTC) stained brain slices of (a) control, (b) TH34, and (c) TH33. Tissue damage was less extensive with cooling, and only a few slices of TH33 had irreversible damage compared to TH34. White tissues indicate irreversible damages (infarcts).

**S9**

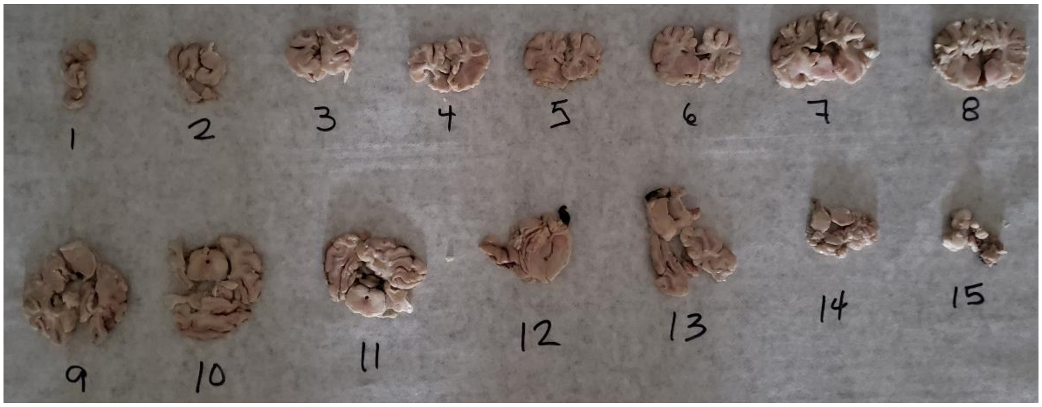


**Fig. S9.** TTC brain slices of an animal euthanized 4 hours after global hypoperfusion was observed in CT perfusion. All the tissues were white, which indicates irreversible damage (infarcts).

**Fig. S2.** Type or paste legend here. Paste figure above the legend.

<insert page break here>

**Table S1.** Type or paste table title here. Paste table below the title.

<insert page break here>

**Table S2.** Type or paste table title here. Paste table below the title.

<insert page break here>

**Movie S1 (separate file).** Type or paste legend here.

**Dataset S1 (separate file).** Type or paste legend here.

**Software S1 (separate file).** Type or paste legend here.
